# Commonalities in alpha and beta neural desynchronizations during prediction in language comprehension and production

**DOI:** 10.1101/2020.05.13.092528

**Authors:** Simone Gastaldon, Giorgio Arcara, Eduardo Navarrete, Francesca Peressotti

**Affiliations:** Dipartimento di Psicologia dello Sviluppo e della Socializzazione (DPSS), University of Padova, Padova, Italy; IRCCS San Camillo Hospital, Venice, Italy

**Keywords:** language prediction, language production, alpha–beta oscillations, internal model

## Abstract

The present study investigates whether predictions during language comprehension are generated by engaging the production system. We recorded EEG from participants performing both a comprehension and a production task in two separate blocks. Participants listened to high and low constraint incomplete sentences and were asked either to name a picture to complete it (production) or to simply listen to the final word (comprehension). We found that in a silent gap before the final stimulus, predictable stimuli elicited alpha (8-10 Hz) and beta (13-30 Hz) desynchronization in both tasks. Source estimation highlighted not only the involvement of the left-lateralized language network, but also of temporo-parietal areas in the right hemisphere. Furthermore, correlations between the desynchronizations in comprehension and production showed spatiotemporal commonalities in language-relevant areas in the left hemisphere, especially in the temporal, lateral inferior and dorsal frontal, and inferior parietal corteces. As proposed by prediction-by-production models, our results show that comprehenders engage the production system while predicting upcoming words.

## 1. Introduction

Top-down prediction of upcoming stimuli has been proposed as a prominent feature of human cognition in order to optimize processing (Clark, 2013; de Lange, Heilbron, and Kok, 2018; Friston, 2005). This has been put forward also for language comprehension, whereby sentential and contextual information guide the preactivation of linguistic representations before it is actually encountered, thus facilitating subsequent elaboration (Federmeier, 2007; Kuperberg & Jaeger, 2016). Prediction has been investigated by employing different paradigms and techniques (see e.g. reading and eye-tracking: Staub, 2005 for a review; visual world paradigm: Huettig, Rommers & Meyer, 2011 for a review; event-related potentials (ERP): Nieuwland et al., 2020 for a large-scale study; Van Petten & Luka, 2012 for a review).

Despite the general agreement on the importance of prediction in language comprehension, what are the linguistic representations involved, the underlying mechanisms and their neural underpinnings is still largely unknown (Huettig, 2015). In the present study we investigated the hypothesis that prediction is implemented by engaging the language production system. To do so, we compared how the same person predicted a target word in two contexts: when s/he had to produce it and when s/he had to listen to it. In order to tap predictive processes, we analyzed the EEG oscillatory activity immediately before the production or the presentation of the target words in contexts in which the target word was either predictable or not. We anticipate that the results revealed large commonalities between predictive processes in the two modalities.

### 1.1 Prediction–by–production models

Traditionally, language comprehension and production have been independently investigated. However, recent work highlights several commonalities in the representations, processes and the underlying neural circuitry (AbdulSabur et al., 2014; Dell & Chang, 2014; Okada & Hickok, 2006; Gambi & Pickering, 2017; Pickering & Garrod, 2014; Silbert, Honey, Simony, Poeppel, and Hasson, 2014). In particular, it has been proposed that prediction during comprehension is implemented through processes traditionally attributed to language production (Huettig, 2015; Pickering & Gambi, 2018; Pickering & Garrod, 2013). The proposals in the literature, however, are not entirely in agreement regarding which processes and representations are involved.

Pickering and Garrod (2013) [P&G2013] envisaged language production and comprehension as a form of action and action perception respectively. In studies of action control, internal forward models are used to predict sensory consequences and future states (Wolpert, 1997; Wolpert & Flanagan, 2001). Similarly, P&G2013 proposed that forward models are used not only to predict the speaker’s own speech during production (Hickok, 2012; Hickok, Houde & Rong 2011), but also to predict others’ speech during comprehension. P&G2013 posited that comprehenders covertly imitate the speakers’ utterance. The inferred communicative intention is fed into a forward model that predicts aspects of upcoming speech (prediction-by-simulation). Such forward models are “impoverished” representations and are extended to all the linguistic hierarchy (semantics, syntax and phonology). According to this view, predictions are rapidly generated without engaging fully-fledged production representations.

According to Huettig (2015), prediction is based on the interaction between multiple mechanisms activated during comprehension (i.e. PACS: production-, association-, combinatorial-, simulation-based prediction). Comprehenders make use of fully-fledged production representations that can be pre-activated through simple associative learning (priming) and through active event simulation. Activation within linguistic representation is further constrained by combinatorial mechanisms sensitive to different linguistic levels. Critically, these mechanisms are shared between comprehension and production.

More recently, Pickering and Gambi (2018) [P&G2018] proposed that the communicative intention derived by the integration of contextual and shared knowledge is fed into the production implementer, as in the PACS model by Huettig (2015). P&G2018 differentiated processes related to prediction-by-association (PA) and to prediction by production (PP). PA is based on the spreading of activation among linguistic levels and it can be equated to semantic/phonological priming. PP is very effective but slow and, since it requires cognitive resources, it is optional. Comprehenders do not necessarily need to go through all the stages of the production implementer and, according to the specific circumstances, they might predict semantic and syntactic features but not the phonology of upcoming words. On the other hand, PA is automatic and mandatory, but less effective. It leads to the pre-activation of all representations that are semantically and phonologically connected, independently of their relevance to the context, which is taken into consideration only in PP.

Summing up, all three proposals assume an important role of priming and event simulation, although for P&G2013 and P&G2018 simulation is part and parcel of the act of production, while in the PACS model it is a separate mechanism interacting with production; P&G2013 ascribe a prominent role to impoverished representations in the form of forward models, while both the PACS model and P&G2018 propose that prediction is based on the implementation of fully-fledged production representations.

### 1.2 Experimental evidence on production-based accounts of prediction

Direct experimental evidence is still relatively scarce. Mani and Huettig (2012) showed that predictive abilities in 2-year old children were correlated with their production vocabulary size (number of words they were able to produce, according to their parents) and not the comprehension vocabulary size (number of words that they could comprehend only). Some ERP studies during sentence reading focused on the N400 effect in relation not only to prediction of meaning but also to prediction of form, showing that form needed longer to pre-activate than meaning (Ito, Corley, Pickering, Martin, and Nieuwland, 2016) and that mismatches at the article preceding the unexpected noun elicited effects at longer latencies for phonology than for grammatical gender (Ito, Gambi, Pickering, Fuellenbach, and Husband, 2020). Assuming that form is encoded after meaning (Levelt, Roelofs, and Meyer, 1999; Indefrey, 2011), these studies suggest that prediction is based on the implementation of production processes. Additionally, it has been shown that taxing the speech production system in a secondary task (silent syllable production) led to reduced N400 responses at the article preceding the unexpected noun, while other secondary tasks (tongue tapping, listening to syllables) did not (Martin, Branzi and Bar, 2018; for a debate on N400 effects at the article, see e.g. Nieuwland et al. 2018; Nicenboim, Vasishth, and Rösler, 2020). Interestingly, a study on German Sign Language (Hosemann, Herrmann, Steinbach, Bornkessel-Schlesewsky, and Schlesewsky, 2013) showed that the N400 effect was already present in the transition between the target unexpected sign and the preceding one, revealing that participants were predicting the phonological features of the upcoming sign, including the trajectory leading from one sign to the other in a modality-specific manner. The authors attribute these modality-specific predictions to forward models, thus supporting a version of production-based accounts of predictions.

Finally, circumstantial evidence is provided by studies on the cerebellum. This structure is contralaterally connected with the neocortex and is assumed to be a crucial node for forward modeling in action and cognition (Ishikawa, Tomatsu, Izawa, and Kakei, 2016; Sokolov, Miall & Ivry, 2017), including language production (Runnqvist et al., 2016; Tourville & Guenther, 2011). Interestingly, the cerebellum has been found to be involved in prediction during comprehension (see Argyropoulos, 2016 and Moberget & Ivry, 2016 for reviews).

Studies of brain-damaged patients present cases of dissociation between comprehension and production abilities, suggesting functional independence between the two systems. However, it should be noted that aphasic deficits in production and comprehension are always a matter of degree rather than all-or-nothing phenomena (Kemmerer, 2015). For instance, Broca’s aphasia, sometimes referred to as ‘agrammatic aphasia’ and traditionally categorized as an expressive impairment, also involves to some degree degraded comprehension (Choy & Thompson, 2010; Rogalsky & Hickok, 2011; Swaab, Brown, and Hagoort, 1997). Interestingly, it has been shown that patients with agrammatic aphasia display impaired lexical prediction (Mack, Ji, and Thompson, 2006). In conclusion, experimental evidence suggests that even though production and comprehension do not fully overlap, the degree of overlap is greater than previously thought.

### 1.3 Neural oscillations in language prediction and production

Differently from ERPs that allow to retain information that is both time- and phase-locked to the onset of a stimulus, time-frequency analysis of the electroencephalographic (EEG) signal enables to observe also the modulation unfolding over time of non-phase-locked oscillatory activity at specific frequency bands (Bastiaansen, Mazaheri & Jensen, 2011). The literature on neural oscillations in language comprehension and production has recently revealed oscillatory correlates of linguistic processing (for reviews, see Meyer, 2018 for speech perception and language comprehension, and Piai and Zheng, 2019 for language production). With respect to the prediction process, the literature is still largely developing. Lewis, Wang and Bastiaansen (2015) proposed that oscillations in the beta band (13-30 Hz) could reflect the generation of predictions whereas oscillations in the gamma band (>30 Hz) could reflect the propagation of prediction error, in line with the domain-general framework proposed by Engel and Fries (2010). From this perspective, in the domain of language processing it has been hypothesized that neural synchronization (reflected in power increase) and desynchronization (reflected in power decrease or suppression) in the beta band signal the maintenance and the change of the current interpretation of the meaning of the sentence. Past literature focused on modulations following syntactic and semantic violations (post-target modulations). These studies observed beta desynchronization in the violation condition (Davidson & Indefrey, 2007; Luo, Zhang, Feng, and Zhou, 2010; Wang et al., 2012), consistent with the idea that sentence structure and meaning, built on the basis of previous context, has to be changed according to the new unexpected target. However, when focusing on pre-target activity, a different modulatory pattern should be observed. In this case during the presentation of the constraining context before the predictable target, the interpretation of the sentence should be changed accordingly, leading to the pre-activation of plausible linguistic information. This change should be associated to the desynchronization of beta band oscillatory activity. Hence, relative to pre- and post-target activity, it can be hypothesized that synchronization (power increase) should be observed after a predictable target and desynchronization (power decrease) should be observed during the interval preceding such a target.

Congruently, oscillatory studies of prediction during comprehension consistently show desynchronization in the beta (but also alpha) range before a predictable target (see Table 1). These studies employed the written modality, with words presented one at a time for fixed durations. While most studies employed high and low constraining sentences (Rommers, Dickson, Norton, Wlotko, and Federmeier, 2017; Wang, Hagoort, and Jensen, 2018), Terporten, Schoffelen, Dai, Hagoort, and Kösem (2020) studied the oscillatory activity pre- and post-target, and the evoked response post-target (M/N400) while reading low, medium and high constraining sentences. The results showed alpha and beta desynchronization before target onset. Interestingly, the oscillatory data showed a non-monotonic relation with constraint level (i.e. the strongest desynchronization was elicited by the medium constraint, followed by the high and then the low constraint). The authors argued that pre-target power modulations reflected working memory demands for target pre-selection. These were maximal for the condition of intermediate levels of constraint in which the pool of activated lexical candidates is larger than in the high constrain condition in which only one candidate is activated. In other studies, however, maintenance in working memory has been more often associated to alpha–beta synchronization (see Weiss & Müller, 2012; Meyer, 2018; Piai, Roelofs, Rommers, Dahlslätt, and Maris, 2015). Moreover, as can be seen in Table 1, effects in oscillatory activity have been detected only in partially overlapping cortical areas. Given these inconsistencies in the results, it is still largely unclear what the processes associated to alpha–beta desynchronization are.^1^

**Table 1:**
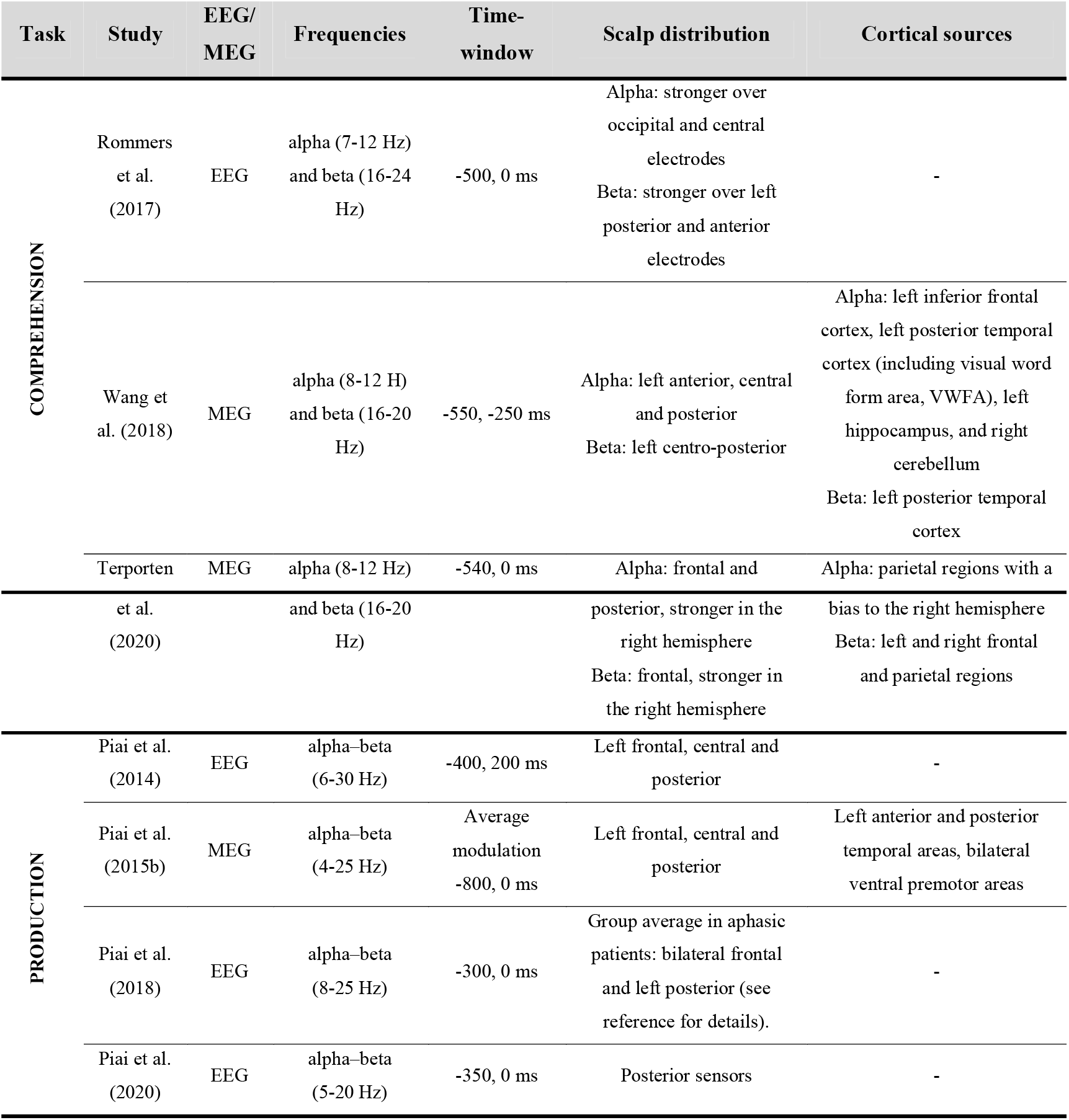
Summary of the studies on neural oscillations pre-target in prediction during comprehension and in context-induced word production. All these studies report desynchronization in the frequency bands and time-windows specified in the table. (EEG: electroencephalography, MEG: magnetoencephalography)

| Task | Study | EEG/<br>MEG | Frequencies | Time-window | Scalp distribution | Cortical sources |
| --- | --- | --- | --- | --- | --- | --- |
| COMPREHENSION | Rommers et al. (2017) | EEG | alpha (7-12 Hz) and beta (16-24 Hz) | -500, 0 ms | Alpha: stronger over occipital and central electrodes<br>Beta: stronger over left posterior and anterior electrodes | - |
|  | Wang et al. (2018) | MEG | alpha (8-12 Hz) and beta (16-20 Hz) | -550, -250 ms | Alpha: left anterior, central and posterior<br>Beta: left centro-posterior | Alpha: left inferior frontal cortex, left posterior temporal cortex (including visual word form area, VWFA), left hippocampus, and right cerebellum<br>Beta: left posterior temporal cortex |
|  | Terporten | MEG | alpha (8-12 Hz) | -540, 0 ms | Alpha: frontal and | Alpha: parietal regions with a |
|  | et al.<br>(2020) |  | and beta (16-20<br>Hz) |  | posterior, stronger in the<br>right hemisphere<br>Beta: frontal, stronger in<br>the right hemisphere | bias to the right hemisphere<br>Beta: left and right frontal<br>and parietal regions |
| PRODUCTION | Piai et al.<br>(2014) | EEG | alpha-beta<br>(6-30 Hz) | -400, 200 ms | Left frontal, central and<br>posterior | - |
|  | Piai et al.<br>(2015b) | MEG | alpha-beta<br>(4-25 Hz) | Average<br>modulation<br>-800, 0 ms | Left frontal, central and<br>posterior | Left anterior and posterior<br>temporal areas, bilateral<br>ventral premotor areas |
|  | Piai et al.<br>(2018) | EEG | alpha-beta<br>(8-25 Hz) | -300, 0 ms | Group average in aphasic<br>patients: bilateral frontal<br>and left posterior (see<br>reference for details). | - |
|  | Piai et al.<br>(2020) | EEG | alpha-beta<br>(5-20 Hz) | -350, 0 ms | Posterior sensors | - |

With respect to language production, in a series of studies, Piai and collaborators focused on the oscillatory correlates of word production by employing context-induced picture naming tasks. In these paradigms, the sentential context preceding the presentation of the target picture either allows or not for predicting the name of the target picture. Time-frequency analyses focus on the interval preceding the target, revealing alpha–beta desynchronization before predictable pictures (see Table 1). An open question is what kind of processes and representations are reflected in the alpha–beta desynchronization found in this kind of production task. Piai, Roelofs, Rommers, and Maris (2015) dissociated the memory- and motor-related components by comparing pre-target beta and alpha desynchronization in two different tasks. In one case the task required to name the picture that followed a constraining or non–constraining sentence frame, in the other case participants were asked to judge whether the picture was predictable or not by pressing a key with their left hand. Results showed alpha–beta desynchronization in different areas, depending on the task. The activity in the left temporal areas and in ventral premotor areas observed during picture naming was associated to word retrieval and speech motor programming. The activity in left posterior temporal and inferior parietal areas and in the right motor area observed during the categorization task were associated to conceptual processing and manual response preparation. In Piai, Klaus and Rossetto (2020), auditory distractors were introduced before picture onset. Alpha–beta desynchronization was delayed when the distractors were semantically related to the target picture with respect to unrelated distractors, suggesting that these power modulations are sensitive to lexico-semantic processing. Along the same lines, Piai, Rommers and Knight (2018) showed that aphasic patients with concomitant left temporal and inferior parietal lesions did not benefit from constraining contexts and did not display the characteristic alpha–beta desynchronization, while patients with left frontal and left temporal (but not inferior parietal) lesions did. According to the authors, this pattern suggests that the desynchronization in the alpha and beta bands elicited in context-induced word production is functionally associated to core semantic memory and lexical retrieval. Whether phonological encoding is captured and reflected in these modulations remains unanswered.

The oscillatory activity in the beta band reported both in prediction during comprehension and in production has led to the hypothesis of a common mechanism shared by the two processes (Molinaro, Monsalve and Lizarazu, 2016). Until now, however, no study has directly compared the oscillatory alpha–beta activity in the two domains. Indirect support pointing towards common mechanisms comes from Pérez, Carreiras, and Duñabeitia (2017) who performed an experiment with hyperscanning where the EEG activity was registered while two participants interacted in a conversation. The results showed that alpha and beta bands oscillations of the speaker and the listener were temporally synchronized. Synchronization within these bands has been interpreted as reflecting coordination between speaker and listener and predictive processing.

### 1.4 The present study

We implemented a within-subject design in which the same participant accomplished a production and a comprehension task in order to directly compare how linguistic information is anticipated in the two tasks. To that end, we targeted the alpha and beta oscillatory activity in an interval immediately preceding the relevant target. More precisely, we used both the cloze probability comprehension task and the context-induced picture naming task in two separated blocks. Participants listened to sentence frames which could either constrain or not towards a target word (see Table 2). After a silent pause of 800 ms, they either listened to the target word or they completed the sentence by naming the target picture. Time-frequency analyses focused on the silent interval between the sentence frame and the target. The structure of the paradigm allowed to directly compare the effects elicited by the same stimuli in the two tasks. In the constraining condition participants could anticipate (and in the production task even plan the response) the target word before hearing it or seeing the corresponding picture. Measuring oscillatory activity before target presentation in the production task allowed us to tap into processes associated to word production planning. The comparison with word prediction during comprehension in the same time interval would highlight the extent to which the two tasks share common mechanisms.

**Table 2:**
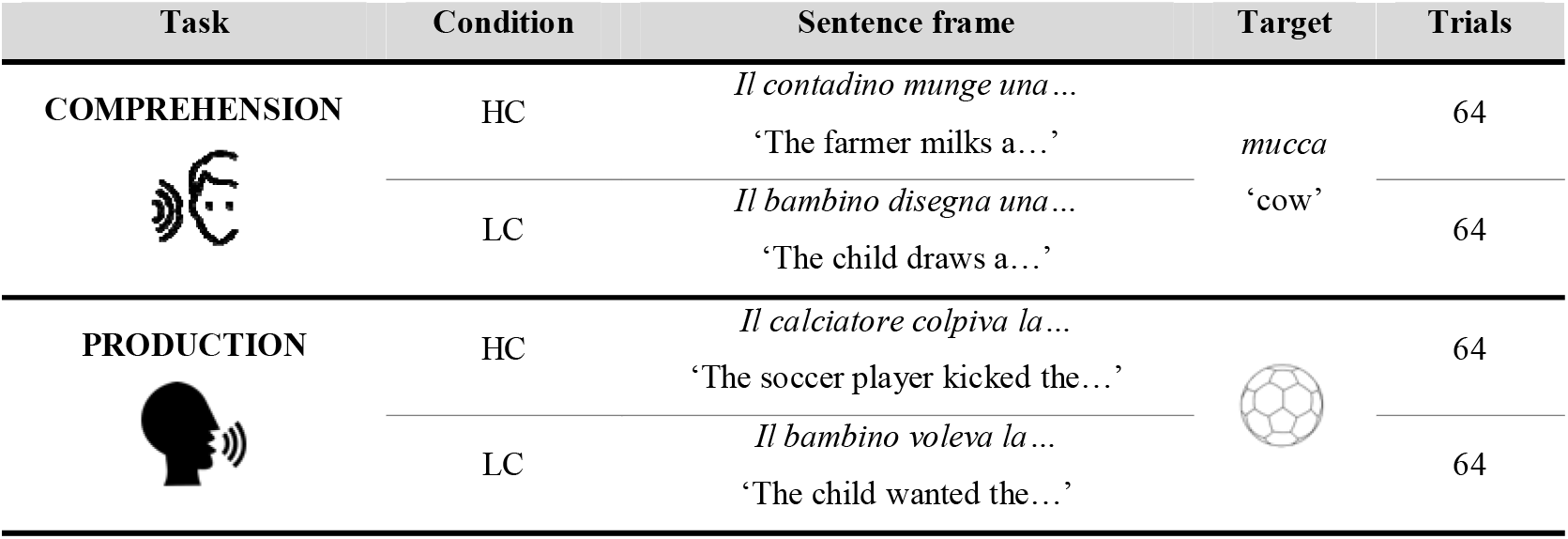
Examples of stimuli used in the experiments.

| Task | Condition | Sentence frame | Target | Trials |
| --- | --- | --- | --- | --- |
| **COMPREHENSION** | HC | *Il contadino munge una...*<br>'The farmer milks a...' | *mucca*<br>'cow' | 64 |
|  | LC | *Il bambino disegna una...*<br>'The child draws a...' |  | 64 |
| **PRODUCTION** | HC | *Il calciatore colpiva la...*<br>'The soccer player kicked the...' |  | 64 |
|  | LC | *Il bambino voleva la...*<br>'The child wanted the...' |  | 64 |

To our knowledge, this is the first study allowing for such direct comparison. In fact, as previously mentioned, shared mechanisms have been proposed in the literature on the bases of similar oscillatory patterns in separate studies investigating either prediction during comprehension or production. In addition, the present study made use of naturalistic auditory stimuli, contrary to most of the previous studies which employed the written modality in an artificial (word-by-word) fashion.

Following the literature, we expect to replicate the pre-target predictability effects of alpha and beta desynchronization in both comprehension and production. If prediction and production share some common mechanisms, we should observe temporal overlaps of alpha–beta modulations between the two tasks in language-relevant areas of the left hemisphere. Moreover, since derivation of communicative intention and event simulation seem essential for anticipating linguistic content both in production and in comprehension, alpha and beta desynchronization could be observed also in cortical regions supporting supramodal integration and traditionally not involved in linguistic processing.

## 2. Materials and methods

### 2.1 Participants

Forty participants were recruited on a voluntary basis (11 males; mean age = 23.7, sd = 4.84). All participants were right-handed native speakers of Italian (handedness evaluated by means of an Italian translation of the Edinburgh Handedness Questionnaire, Oldfield, 1971; mean laterality index = 86, sd = 15.28). None of them reported a history of neurological, language-related or psychiatric disorders. All participants signed an informed consent to participate in the experiment. The study was approved by the Ethical Committee for the Psychological Research of the University of Padova (protocol n. 2920).

### 2.2. Stimuli

One hundred twenty-eight concrete, animate and inanimate nouns were selected and paired with a black-and-white line picture (240 x 240 pixels) representing the word referent. For each picture, a scrambled version was also created, in such a way that the referent was not recognizable. For each target noun, two sentence frames were constructed: one whose semantic content leads to the target word with a high probability (highly constraining; HC) and one for which the target word is not particularly likely but is still plausible given the sentential content (low constraining; LC, see Table 2). This resulted in 256 sentences in total (128 HC, 128 LC). Sentence frames associated to the same target were matched for number of syllables, had a similar syntactic structure, and had the same article or preposition as the final word. The constraint was modeled as cloze probability (CP) of the target word given the frame, assessed with an online sentence completion questionnaire involving 71 respondents, none of whom took part in the subsequent experiment, who were asked to complete each sentence frame with the word they considered most appropriate (HC sentences: mean CP = 0.873, sd = 0.092; LC sentences: mean CP = 0.052, sd = 0.078). Subsequently, all sentence frames and target words were recorded from a female native speaker in a quiet room using a microphone connected to a PC using Audacity (sampling rate of 44.1 KHz). Frames and targets were recorded separately. The speaker was instructed to keep the reading pace as steady as possible and to keep a constant distance from the microphone. Sentence frames were then appropriately trimmed at the beginning and at the end using Audacity. The approximate number of syllables per second, assuming a constant pace, for each sentence frame was estimated as the number of syllables of the sentence divided by the length of each audio file.

Target words and their associated sentence frames were then divided into two lists, A and B, each containing 64 target words and the associated 128 sentence frames (64 HC and 64 LC). The two lists were matched for lexical frequency (log-scaled; obtained from *COLFIS*, Bertinetto et al., 2005), number of phonemes and number of syllables (obtained from *PhonIta 1.10*, Goslin, Galluzzi, and Romani, 2014), number of syllables per second, audio file duration, both across conditions and within conditions. The difference of CP was not significant across condition, and was significant between conditions, both in the whole set and within each list (see Table 3 for stimuli matching).

**Table 3:** Variables controlled across lists and conditions (Welch’s *t*-tests). Means and standard deviations (in parenthesis) are reported. HC: high constraint, LC: low constraint.

| LISTS |  |  |  |  |  |
| --- | --- | --- | --- | --- | --- |
|  | Mean (sd) List A | Mean (sd) List B | *t*-value | df | *p*-value |
| Lexical frequency (log-scaled) | 3.776 (1.251) | 3.847 (1.31) | -0.51 | 235.75 | 0.6104 |
| No. phonemes (word) | 6.063 (1.701) | 6.281 (1.631) | -1.05 | 253.55 | 0.2947 |
| No. syllables (word) | 2.578 (0.728) | 2.672 (0.641) | -1.09 | 250.06 | 0.2751 |
| No. syllables (sentence frame) | 10.680 (2.012) | 10.672 (2.248) | 0.03 | 251.07 | 0.9766 |
| Audio length (sec) (sentence frame) | 2.388 (0.397) | 2.381 (0.414) | 0.15 | 253.54 | 0.8851 |
| No. syllables/sec (sentence frame) | 4.482 (0.485) | 4.480 (0.523) | 0.05 | 252.52 | 0.9643 |

| **Cloze probability overall** | 0.456 (0.421) | 0.468 (0.421) | -0.23 | 245 | 0.8215 |
| --- | --- | --- | --- | --- | --- |
| **Cloze probability HC** | 0.868 (0.094) | 0.878 (0.09) | -0.62 | 125.84 | 0.5351 |
| **Cloze probability LC** | 0.045 (0.068) | 0.058 (0.086) | -1 | 119.69 | 0.3193 |
| **CONDITIONS** |  |  |  |  |  |
|  | **Mean (sd) HC** | **Mean (sd) LC** | ***t*-value** | **df** | ***p*-value** |
| **No. syllables (sentence frame)** | 10.750 (2.074) | 10.602 (2.182) | 0.56 | 253.34 | 0.5774 |
| **Audio length (sec) (sentence frame)** | 2.413 (0.407) | 2.356 (0.402) | 1.12 | 253.97 | 0.2629 |
| **No. syllables/sec** | 4.459 (0.441) | 4.504 (0.56) | -0.71 | 240.87 | 0.4758 |
| **Cloze probability overall** | 0.873 (0.092) | 0.052 (0.077) | 77.47 | 246.66 | <0.001 |
| **Cloze probability List A** | 0.868 (0.094) | 0.045 (0.068) | 56.96 | 114.71 | <0.001 |
| **Cloze probability List B** | 0.878 (0.09) | 0.058 (0.086) | 52.69 | 125.62 | <0.001 |

### 2.3 Procedure

Participants were seated in a comfortable chair in a soundproof room with a computer connected to a CRT monitor, built-in speakers, a keyboard and a microphone to record responses. Stimuli were presented with E-Prime 2.0 (Psychology Software Tools, Pittsburgh, PA). Each participant performed the comprehension task and the production task in a blocked design. The structure of the trials in the two tasks is shown in Figure 1. After a silent interval of 800 ms, a sentence frame was played through the computer speakers, and it was followed by a second silent gap of 800 ms. Throughout this phase the fixation cross remained on the screen. Afterwards, the target was presented together with a visual stimulus for 2 seconds. In the comprehension task the visual stimulus was constructed by scrambling the picture corresponding to the target in such a way that the referent was not recognizable. In the production task, the visual stimulus was the picture of the target word. In the comprehension task, participants were instructed to listen carefully to the sentence. To ensure that they paid attention to the sentence, 26 trials (20%) included a statement about the preceding sentence appearing as written text after the target for 2 seconds. Participants were asked to judge whether it was true or false by providing a vocal response. In the production task, participants were instructed to name the picture as fast and as accurately as possible.

**Figure 1:**
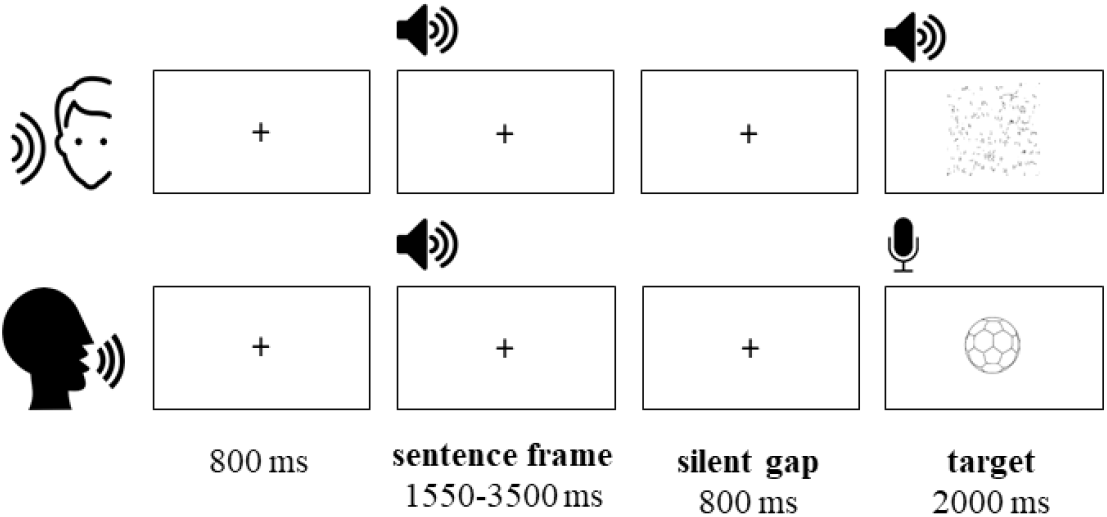
Trial structure in the comprehension (top) and the production (bottom) tasks.

For each participant, list A or B was associated to one of the tasks (e.g. list A for comprehension and list B for production). Task order and the lists associated to the tasks were counterbalanced across participants, resulting in a 2×2 design (2 lists × 2 tasks). Trial order presentation was pseudo-randomized for each participant by using Mix (van Casteren & Davis, 2006) in such a way that the minimum number of trials between the first and the second presentation of the same target word was seven, and no more than three consecutive trials belonged to the same condition. The inter-trial interval varied from trial to trial (1, 1.2 and 1.5 seconds). After every 32 trials participants could take a short break. Responses were recorded through the microphone, positioned at a fixed distance from the participant (~50 cm). During the experimental session participants were instructed to minimize eye movements, blinks and facial muscle activity during the presentation of the stimuli. Before each task, a training session of 8 trials (not included in the experimental session) was used to familiarize the participant. Each task lasted approximately 20 minutes.

### 2.4 Response coding and production RT analyses

For the comprehension task, true/false responses were coded as correct or incorrect. Trials with incorrect responses were excluded from the EEG analyses.

In the production task, audio recording started at the onset of the picture and lasted for 2 sec. Responses were manually coded as incorrect when participant 1) failed to provide an answer, 2) produced hesitation sounds, 3) started producing a word but then produced another word, 4) produced the correct target word before recording onset. Trials with incorrect responses were excluded from the EEG analyses. Response onset was measured from each audio recording using Chronset (Roux, Armstrong, and Carreiras, 2017). In case Chronset returned some *NA* values, the correspondent audio waveforms were inspected manually with Audacity in order to determine the response onset. The set of correct responses was then analyzed using R (R Core Team, 2014).

RTs were analyzed by means of linear mixed-effects models (Baayen, Davidson, and Bates, 2008) using the *lme4* package (Bates, Mächler, and Walker, 2015), with random intercept for participant and target word. The *lmerTest* package (Kuznetsova, Brockhoff, and Christensen, 2017) was used to estimate the *p*-values for model parameters. First, a null model including random effects was computed, and in each subsequent model a predictor or an interaction between predictors was added. An ANOVA between models was then performed, and the best-fit model was selected considering AIC (Akaike Information Criterion) and BIC (Bayesian Information Criterion) as indices of fit and the *p*-value of the test between models.

### 2.5 EEG data acquisition and pre-processing

Electroencephalogram was recorded with a system of 64 active Ag/AgCl electrodes (Brain Products), placed according to the 10–20 convention (ActiCap). Sixty of them were used as active electrodes (Fp1, Fp2, AF3, AF4, AF7, AF8, F1, F2, F3, F4, F5, F6, F7, F8, Fz, FT7, FT8, F1, F2, F3, F4, F5, F6, Fz, FC1, FC2, FC3, FC4, FC5, FC6, T7, T8, C1, C2, C3, C4, C5, C6, Cz, TP7, TP8, CP, CP2, CP3, CP4, CP5, CP6, CPz, P1, P2, P3, P4, P5, P6, P7, P8, PO3, PO4, PO7, PO8, PO9, PO10 POz, O1, O2, Oz). Reference was placed at the left earlobe. Three electrodes were used to record blinks and saccades (external canthi and below the left eye). Electrode impedance was kept below 10 kΩ throughout the experiment. The signal was amplified and digitized at a sampling rate of 1000 Hz. Before the tasks, a resting state of 5 minutes was recorded, which is not analyzed further here. Each task was recorded separately. As a result, 3 recordings were obtained for each participant (resting state, production, comprehension).

Pre-processing and analyses were performed using the MATLAB toolbox Brainstorm (Tadel, Baillet, Mosher, Pantazis, and Leahy, 2011; Tadel et al. 2019), which is documented and freely available for download online under the GNU general public license. A high-pass filter at 0.5 Hz with 60 dB attenuation was applied to the raw data. Noisy or flat channels were marked as ‘bad’ and excluded (max 2 channels marked as ‘bad’ per participant). No interpolation of bad channels was performed. Segments with extreme muscle artifacts were marked as ‘bad’. Subsequently, Independent Component Analyses (ICA) with 60 components was computed to detect and remove artifact components with known time-series and topographies (blinks, saccades, and power-line noise at 50 Hz).^2^ Markers for incorrect responses were manually added to the continuous EEG recording according to the off-line evaluation of the audio files. Finally, 3-second epochs (from −1.5 to 1.5 s) were imported around two event markers: (1) the onset of the trial (fixation cross), and (2) the onset of the 800 ms gap pre-target. The epochs in (1) were not divided into conditions and constitute the condition-average baseline for the event-related synchronization / desynchronization (baseline_comp_ and baseline_prod_). This ensures a higher signal-to-noise ratio given the higher number of trials included as baseline, and therefore a better estimate of the relative power change (Cohen, 2014). The epochs in (2) were divided into HC and LC conditions (HC_comp_, LC_comp_, HC_prod_ and LC_prod_). All epochs were visually inspected, and those with artifacts (uncorrected blinks/saccades, muscle activity, channel drifts, transient electrode displacements) were rejected. All trials in (2) which included a marker of incorrect response were rejected.

### 2.6 Time-frequency decomposition and statistical analyses (sensor-level)

In the time-frequency (TF) decomposition, power was computed by using Morlet wavelets. According to Morlet Wavelet implementation in Brainstorm software, wavelets were built starting from a mother wavelet with central frequency = 1 and FWHM = 3 (7-cycle wavelets), and then generating new wavelets spanning from 5 Hz to 30 Hz with step 1 Hz.^3^ TF maps were obtained for each trial for all conditions ^(baseline^comp^, HC^comp^, LC^comp^, baseline^prod^, HC^prod^, LC^prod^)^. ^Due to the large windows for^ epoching (3 seconds), edge effects at the selected frequencies did not involve the windows of interest. Subsequently, TF maps were averaged within each condition for each participant.

Event-related synchronization/desynchronization (ERS/ERD) was used as normalization method.^4^ For each participant, the average TF map of the two conditions were normalized against the mean computed over the interval [-550 −250] ms of the average baseline TF map (baseline_comp_ for HC_comp_ and LC_comp_; baseline_prod_ for HC_prod_ and LC_prod_). This yielded the %-change of power over time relative to the baseline for each frequency.

After having obtained the normalized TF map for each participant, non-parametric cluster-based permutation tests were performed for each task on the 800 ms pre-target gap (paired one-tailed *t*-test with cluster correction; Maris & Oostenveld, 2007). The critical α level was set to 0.05, the minimum number of neighboring channels set to 2, and the number of Monte Carlo simulations for the permutations to 1000. Following existing literature, we used one-tailed tests to ensure higher statistical power to detect an effect in a specific direction. Specifically, the expected alternative hypothesis was that HC conditions elicited reduced power compared to LC conditions. An additional two-tailed analysis with cluster-based permutation (same parameters as above) was performed also between the differentials (HC–LC) between tasks (comprehension *vs* production) to test for an interaction. From now on we refer to the difference between HC and LC in each task as Δ_comp_ and Δ_prod_, and to the statistical contrast between them as interaction.

### 2.7 Time-frequency decomposition and statistical analyses (source-level)

To estimate EEG activity at source level we implemented the following steps. First, a noise covariance matrix for each task was computed from the baseline epochs in the time-window [-550 −250] ms. OpenMEEG BEM (Boundary Element Method) with 8002 vertices was used as forward solution^5^ (Gramfort, Papadopoulo, Olivi, and Clerc, 2010) with ICBM152 as template anatomy. This method models three realistic layers (scalp, inner and outer skull) in addition to the cortical surface; for this reason, it is recommended for EEG data, given the differential electrical propagation through the different types of tissue. Minimum Norm Imaging (NMI) normalization with sLORETA (Standardized Low Resolution Brain Electromagnetic Tomography; Pascual-Marqui, Michel, and Lehmann, 1994) was used as inverse solution. The dipole orientation was unconstrained, to obtain a better estimation in lack of individual anatomy scans.^6^ Timefrequency decomposition was performed on each epoch, averaged and normalized against the baseline as for the TF at sensor level. TF maps were then averaged across frequencies in four bands: alpha (8-12 Hz), beta1 (13-19 Hz), beta2 (20-25 Hz) and beta3 (26-30 Hz). Subsequently, ERS/ERD maps were downsampled at 150 Hz, to reduce the computational burden. Cluster-based permutation tests (one-tailed paired *t*-tests) for main effects and interaction (two-tailed paired *t*-tests) on source-space TF data were performed as previously described.

### 2.8 Between-task source-level correlations

Pearson correlations between Δ_comp_ and Δ_prod_ at the source level were performed. This provides an estimate of putative shared cortical generators of the desynchronizations in prediction during comprehension and in word planning in production. Correlations were computed separately for the alpha band (8-12 Hz) and the three beta sub-bands (13-19, 20-25 and 26-30 Hz) on %-power change averaged in intervals of 200 ms (0-200, 200-400, 400-600 and 600-800 ms), resulting in 16 correlation maps. For each vertex of the cortex model, two vectors of values were correlated. Each vector contained 36 values, one for each participant, representing the average Δ%-power change at a given frequency band and time-window in the two tasks. We decided to average in time because it is likely that cortical modulations underlying possible shared processes are not temporally aligned across the two tasks due to different demands influencing participants’ performance. In this way we can capture desynchronizations at the same vertex that are slightly shifted in time. For each frequency band, correlations were thresholded for *p* < 0.05 and minimum size = 50 (number of connected vertices), in order to exclude not only statistical non-significant correlations, but also statistically significant but isolated and likely meaningless correlations, given the very low spatial resolution of the technique. Then, correlation maps were inspected, and the interval with the strongest and more spatially extended correlations were identified. In a more exploratory fashion, we performed additional correlations on the averages of the identified time-windows in order to provide a clearer summary of the results and capture the possible commonalities between the tasks. For this additional analysis, we only focused on the positive correlations.

## 3. Results

### 3.1 Word production response times

Response accuracy was very high (98.5%). Only 84 responses were coded as incorrect, 59 in the LC condition and 25 in the HC condition. Error rates were not analyzed. Figure 2 shows response times of correct trials divided by condition.

**Figure 2:**
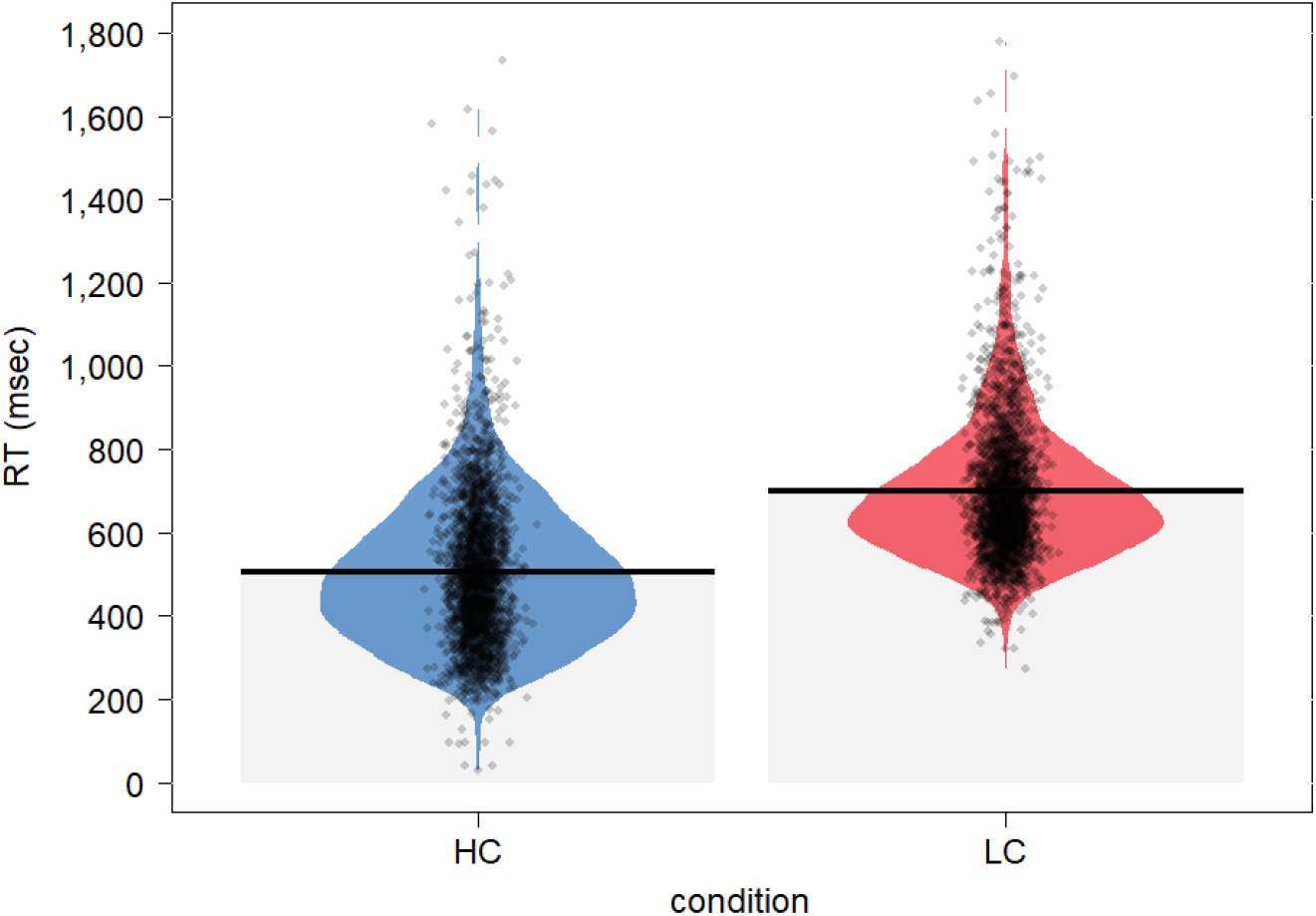
Violin plot of the response times of correct trials in the production task for the HC and the LC conditions. HC: mean = 507 ms, sd = 184.786; LC: mean = 698 ms, sd = 172.22.

Latencies of correct responses were fitted to mixed-effects models; Table 4 shows the results of the ANOVA between models. The model which best explained the data is M4, which included Repetition, Condition, Lexical frequency and the interaction Condition × Lexical Frequency as fixed effects.

**Table 4:** Statistics of model selection.

| Model | Effects | Df | AIC | BIC | X<sup>2</sup> | p-value |
| --- | --- | --- | --- | --- | --- | --- |
| M0 | Random effects (R.E.) | 4 | 66966 | 66992 |  |  |
| M1 | R.E. + Repetition | 5 | 66705 | 66737 | 263.22 | < 0.001 |
| M2 | R.E. + Repetition + Condition | 6 | 64857 | 64897 | 1849.1 | < 0.001 |
| M3 | R.E. + Repetition + Condition + Lexical<br>Frequency | 7 | 64858 | 64904 | 1.134 | 0.287 |
| M4 | R.E. + Repetition + Condition + Lexical<br>Frequency + Condition × Lexical<br>Frequency | 8 | 64851 | 64903 | 9.258 | 0.002 |
| M5 | R.E. + Repetition + Condition + Lexical<br>Frequency + Condition × Lexical<br>Frequency + Condition × Repetition | 9 | 64852 | 64911 | 1.231 | 0.267 |

The model showed a main effect of Repetition (estimate = −85.023, *t* = −20.74, *p* < .001, 95% CI: −93.060 − −76.990) - with estimated faster responses at the second presentation of the same target picture – and of Condition (estimate = 232.135, *t* = 17.909, *p* < .001, 95% CI: 206.724 - 257.545) – with estimated faster responses in the HC relative to LC condition. There was no main effect of Lexical Frequency (*p* = .959), but there was an interaction between Frequency and Condition (estimate = −9.795, *t* = −3.044, *p* < .01, 95% CI: −16.102 − −3.487): the effect of Lexical Frequency was present in the LC condition, with decreasing RTs when lexical frequency increases. Table 5 shows all the parameter estimates of the model.

**Table 5:** Parameter estimates of model M4.

| Parameter | Estimate | SE | df | *t*-value | *p*-value | 95% CIs |
| --- | --- | --- | --- | --- | --- | --- |
| **Intercept** | 547.970 | 22.126 | 188.392 | 14.766 | < 0.001 | [504.384 – 591.563] |
| **Repetition** | -85.023 | 4.099 | 4870.025 | -20.740 | < 0.001 | [-93.060 – -76.990] |
| **Condition** | 232.135 | 12.962 | 4870.763 | 17.909 | < 0.001 | [206.724 – 257.545] |
| **Lexical Frequency** | 0.236 | 4.595 | 162.165 | 0.051 | 0.959 | [-8.822 – 9.297] |
| **Condition × Lexical Frequency** | -9.795 | 3.128 | 4870.514 | -3.044 | < 0.01 | [-16.102 – -3.487] |

### 3.2 Sensor-level time-frequency analysis

The data of two participants were excluded from the analyses due to an excess of trials coded as incorrect in the comprehension task (34.6% and 53.9%). Another two participants were excluded due to excessively noisy recordings in the EEG. The mean percentage of epochs retained are the following: baseline_comp_: 88.8%, HC_comp_: 89.7%, LCcomp: 87.5%, baselineprod: 87.7%, HCprod: 89.3%, LCprod: 89.7%.

The cluster-based permutation tests contrasting HC *vs* LC conditions in the two tasks were significant. In the comprehension task, the effect was associated to a negative cluster (*p* = 0.003, *t*-sum = −324052, size = 134070; Figure 3a). In the production task, the effect was associated to a negative cluster (*p* = 0.001, *t*-sum = - 946882, size = 361719; Figure 3b). This suggests that high word predictability elicited desynchronization before target presentation, be it an auditory word or a picture to name overtly. The effects were widespread across all sensors and appeared to span the entire alpha and beta ranges, with variability of modulations across the gap. The analysis testing for the interaction did not yield significant results (all clusters *p* > 0.05; see Supplementary Material).

**Figure 3:**
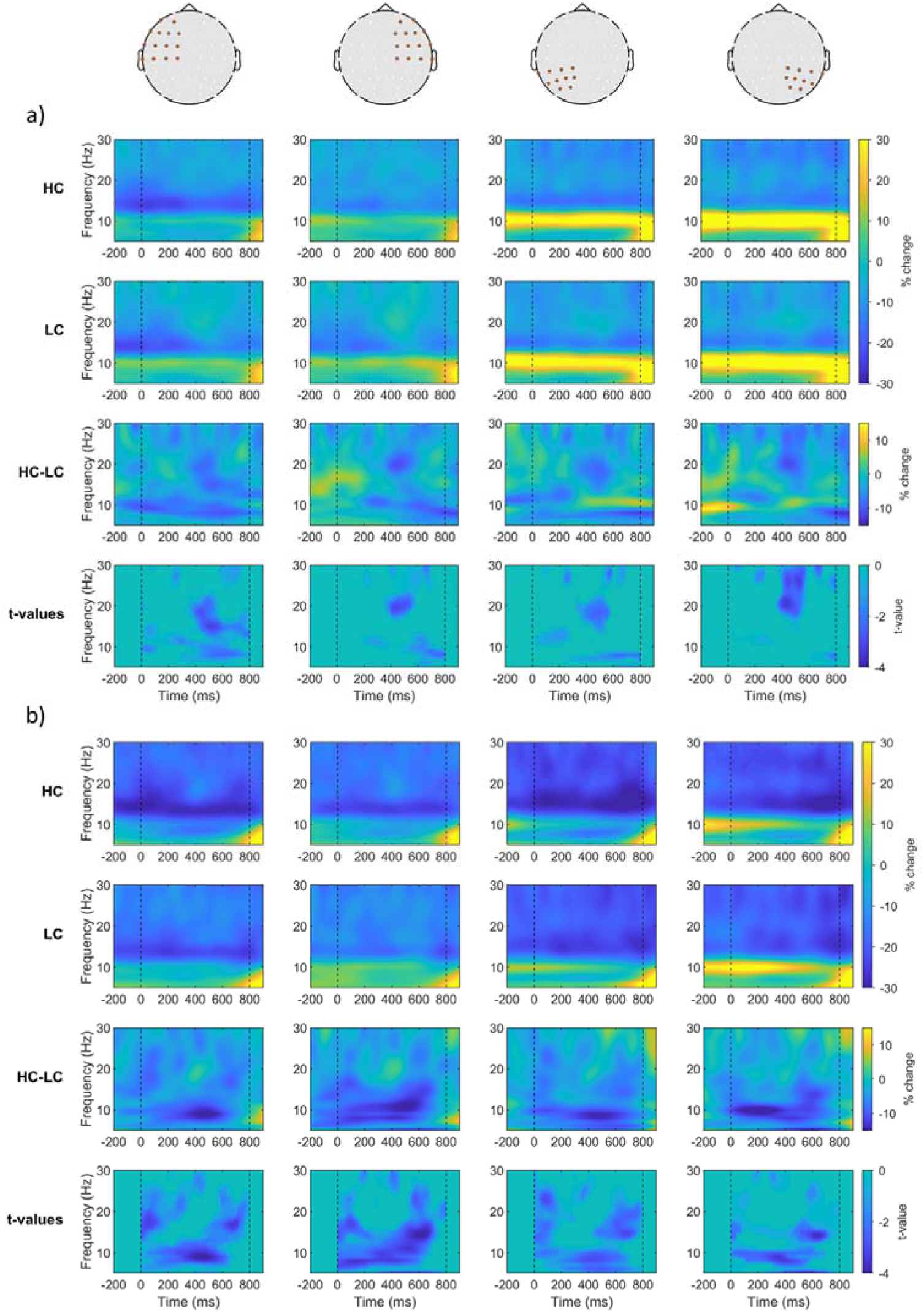
Time-frequency maps averaged across groups of sensors for the comprehension (a) and the production (b) tasks. The averaging is only for visualization purposes; statistical testing was performed on all electrodes. Each column represents a group of sensors, specified on the scalp model above. The rows represent the average TF maps of the HC condition, the LC condition, the HC-LC differential (Δ_comp/prod_), and the *t*-values of the statistical contrast (*t*-maps are masked for values associated to the significant cluster).

### 3.3 Source-level time-frequency analysis

Source-level contrasts identified two significant negative clusters in the comprehension task (cluster 1, left hemisphere: *p* = 0.02597, *t*-sum = −742975, size = 326126; cluster 2, right hemisphere: *p* = 0.04995, *t*-sum = −645503, size = 280809) and two negative clusters in the production task (cluster 1, left hemisphere: *p* = 0.001, *t*-sum = −1820707, size = 719067; cluster 2, right hemisphere: *p* = 0.002, *t*-sum = −1405540, size = 562307). The interaction analysis did not yield significant results (all clusters *p* > 0.05; see Supplementary Material). Results are shown in Figure 4a and Figure 4b.

**Figure 4:**
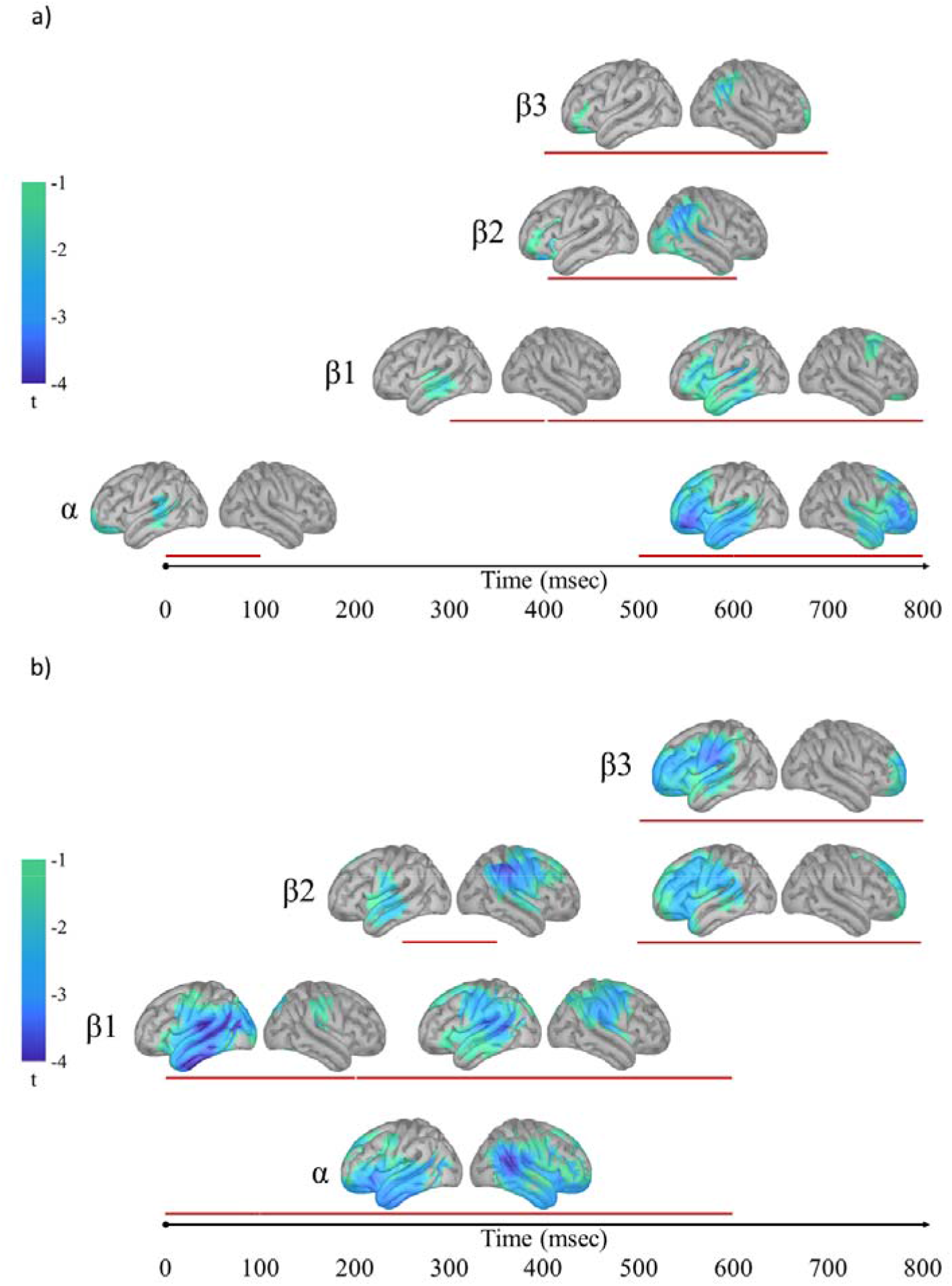
Summary of the statistical contrasts between HC and LC conditions in the comprehension (a) and production (b) tasks. Selected cortical maps of *t*-values are shown for each of the frequency bands (α: 8-12 Hz; β1: 13-19 Hz; β2: 20-25 Hz; β3: 26-30 Hz) in averaged time-windows determined from inspecting the time-course of the results. For each map, the time-window used for averaging is indicated by the red line below each plot (comprehension: α: 0-100, 500-800 ms; β1: 300-400, 400-800 ms; β2: 400-600 ms; β3: 400-700 ms; production: α: 0-600 ms; β1: 0-200, 200-600 ms; β2: 250-350; 500-800 ms; β3: 500-800 ms). Averaging is for visualization purposes only; analyses were performed on all timepoints after downsampling. Only *t*-values ranging from −1 to −4 and part of a significant cluster are shown. Complete results are provided in the Supplementary Material.

In comprehension, alpha desynchronization was stronger towards the end of the gap and involved the bilateral frontal and temporal cortex; early in the gap it involved the left posterior temporal cortex. Beta desynchronization was found in the temporal (beta1) and inferior frontal (beta1, beta2, beta3) corteces of the left hemisphere, and in temporo-parietal-occipital areas (beta1, beta2, beta3) of the right hemisphere. In production, alpha desynchronization involved the bilateral prefrontal, temporal and inferior parietal corteces, with a bias in the right hemisphere. Beta desynchronization involved an extended cortical network, including the temporal, parietal and frontal cortex in the left hemisphere, and the parietal cortex in the right hemisphere.

### 3.4 Source-level correlations

Figure 5 shows positive correlations between Δ_comp_ and Δ_prod_ as defined in the Method section (complete correlation maps are reported in the Supplementary Material). The areas highlighted include the left temporal cortex, the inferior frontal cortex, motor and supplementary motor corteces, the left insula, and the left inferior parietal cortex.

**Figure 5:**
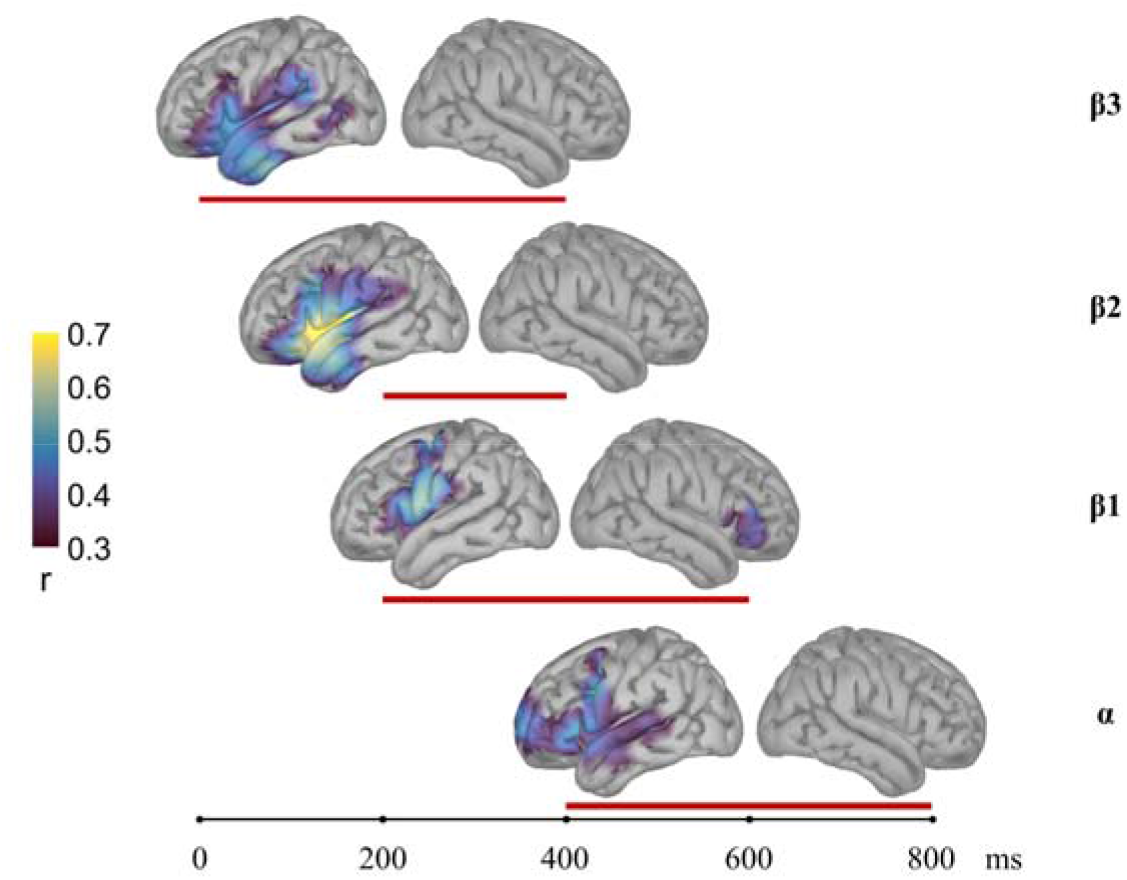
Positive correlations between Δ_comp_ and Δ_prod_ at the source level at each frequency band. The timeline at the bottom represents the 800 ms silent interval between sentence frame and target; the red lines below each cortical map represent the time-window of the correlation displayed above it (α: 400-800 ms; β1: 200-600 ms; β 2: 200-400 ms; β3: 0-400 ms).

## 4. Discussion

We employed a within-subject design in order to directly compare alpha and beta oscillatory modulations elicited by predictive processes in comprehension and production by manipulating cloze probability. We found alpha and beta desynchronization in HC relative to LC conditions preceding the target stimulus. The cortical sources appeared to be left frontal, temporal and inferior parietal, involving the traditional language network, but also right parietal and temporo-parietal, possibly reflecting ‘extra-linguistic’ processes. Positive correlations between Δ_comp_ and Δ_prod_ were found in the left temporal, frontal, and inferior parietal cortices.

In the following paragraphs we will discuss: 1) the desynchronizations in the left hemisphere in relation to language-specific and domain-general processing; 2) the desynchronizations in the rTPJ in relation to contextual updating of internal models; 3) how these results are compatible with production-based accounts of prediction; 4) limitations of the present study and future developments and directions.

### 4.1 Alpha–beta desynchronization in the left language network as index of predictive information retrieval and encoding

In both tasks, HC contexts induce desynchronization of the alpha and beta bands relative to LC contexts. We interpret the desynchronization as marker of preactivation of linguistic information, both in predicting during comprehension and in planning for word production.

In the comprehension task, the language network is engaged in actively updating the sentence-level representation in a top-down fashion. In the HC condition the preceding context allows for the generation of strong predictions about the upcoming word. The information retrieved from long-term memory in this case is rich and specific. This predictive process is reflected in the desynchronization of oscillatory activity in the alpha and beta band observed in language-relevant areas (left temporal and left inferior frontal areas).

In the production task, the HC condition leads to faster naming latencies with respect to the LC condition. Moreover, the effect of lexical frequency was found in the LC but not in the HC context (replicating previous studies, Griffin & Bock, 1998; Piai et al., 2014). The pattern clearly signals that lexical retrieval in the HC condition occurs before picture onset, and the alpha–beta desynchronization effect observed before picture presentation reflects the retrieval of specific lexical information.

These conclusions are compatible with the ‘information by desynchronization’ hypothesis put forward by Hanslmayr, Staudigl, and Fellner (2012). According to this study, information encoding and retrieval is associated with desynchronized firing of neural populations in the alpha and beta frequencies. By applying mathematical modeling, the authors showed that the power of local field potentials at these frequencies (and consequently of the scalp-level EEG fluctuations) is negatively related to the richness of information represented in the brain. The stronger the desynchronization of neural populations, the stronger the decrease in alpha–beta power, and the richer the information encoded or retrieved. From this perspective, therefore, our results are in line with studies showing that alpha–beta desynchronization is related to prediction precision in spatial attention (Bauer, Stenner, Friston, and Dolan, 2014) and pitch change (Chang, Bosnyak, and Trainor, 2018), in successful word memory formation (Griffiths, Mazaheri, Debener, and Hanslmayr, 2016) and word encoding (Meeuwissen, Takashima, Fernández, and Jensen, 2011), and fidelity of stimulus-specific information tracking in the visual and auditory domains (Griffiths et al., 2019).

In conclusion, our findings are in line with previous studies showing alpha–beta desynchronization in the language network before target onset, compatible with the role of alpha–beta desynchronization in signaling active change in the cognitive set.

### 4.2 Alpha–beta desynchronization in the right temporo-parietal junction as index of internal modeling and expectation updating

The alpha and beta desynchronization effects we reported also involved the right hemisphere. Source analyses showed that the effect extended mainly in the prefrontal cortex (alpha range) and in the occipito-temporo-parietal cortex (beta range) in comprehension, and in the temporal, parietal and prefrontal cortex (both alpha and beta ranges) in production. Terporten et al. (2020) reported oscillatory effects in the right parietal cortex in a comprehension task and interpreted this activity as reflecting attentional and working memory demands. The effect we found in the same area in both tasks may capture attentional effects, particularly in the production task which requires participants to shift from listening to preparing to speak.

Our results also suggest that a particular region, the right temporo-parietal junction (rTPJ), is involved in anticipatory language processing in both comprehension and production, although not entirely in the same frequency bands (in beta2-3 in comprehension; alpha and beta1-2 in production). The label ‘rTPJ’ does not describe a well-defined region, and it includes portions of the inferior supramarginal gyrus, the angular gyrus, the posterior superior and middle temporal sulcus and gyrus (Geng & Vossel, 2013). In the literature, this area has been considered to be implicated in different domains of cognition, including bottom-up attention, self-perception, episodic memory, social cognition and Theory of Mind (Igelström & Graziano, 2017). Interestingly, by using fMRI it has been shown that the networks involved in language comprehension and in Theory of Mind – especially the rTPJ, – are synchronized at rest and during story comprehension (Paunov, Blank, and Fedorenko, 2019), implying that coordinated processing of linguistic and social information is crucial for communication. Moreover, activity in the right or bilateral TPJ has been associated to the processing of pragmatic aspects of language (Bašnáková, Weber, Petersson, Van Berkum, and Hagoort, 2014; Carotenuto et al., 2018; Spotorno, Koun, Prado, Van Der Henst, and Noveck, 2012).

Based on the vast literature on rTPJ, Geng and Vossel (2013) proposed a unifying role of rTPJ as a hub whose specific function is determined by the network it is co-activated with. Specifically, the authors suggested that this region is responsible for the contextual updating of internal models in order to adjust expectations about upcoming events and guide planning of future actions. It is likely that during language processing this region contributes to the interfacing of extra-linguistic with linguistic information (including pragmatic knowledge). Our results support this view and suggest that rTPJ works in concert with the left-lateralized language network to construct and update an internal model of the communicative intention for the generation of contextually appropriate predictions and responses. Congruent with prediction-by-production models, the effects in rTPJ highlighted the contribution of contextually appropriate information beyond mere priming (Camblin, Gordon & Swaab, 2007; Hintz, Meyer, and Huettig, 2020; Lowder & Ferreira, 2016; Otten & Van Berkum, 2008).

### 4.3 Compatibility with prediction-by-production models

We observed that in both comprehension and production alpha–beta power decreased before encountering a predictable stimulus. To what extent do these effects reflect common processes involving shared representations? The positive correlations between desynchronizations in the two tasks reveal common activity in areas of the left hemisphere, and more specifically the anterior temporal, inferior parietal, temporo-parietal, and inferior frontal corteces. It is relevant to note that all these areas are generally associated not only with lexical-semantic retrieval but also with word production. This suggests that when predicting a word, comprehenders engage, at least to some extent, the word production network, as proposed by prediction-by-production accounts (Huettig, 2015; Pickering & Gambi, 2018; Pickering & Garrod, 2013). The consensus on the neural bases of word production is that lexical-semantic retrieval involves the anterior and middle temporal cortex, phonological retrieval involves the posterior temporal–inferior parietal cortex and syllabic planning involves the inferior frontal cortex and premotor cortex, which activates the associated articulatory sequences in the motor cortex (for reviews see Indefrey, 2011; Roelofs & Ferreira, 2019; Strijkers & Costa, 2016). Correlation analyses highlight the involvement of all these areas, suggesting that prediction in comprehension implies activation of fully-fledged representations involved in word production.

In the previous section we suggested that the effects found in the rTPJ may reflect the modeling of communicative intention and the updating of such model. Both Huettig (2015) and Pickering and Gambi (2018) propose that this kind of internal modeling is part of the prediction-by-production accounts: comprehenders simulate the communicative intention of the speaker and feed this to the production implementer. We argue that the lack of temporal overlap between task desynchronizations in this region may be due to the fact that the actual intention to produce a word may be responsible for the anticipation of the recruitment of rTPJ in the production task. Indeed, the effect emerged in beta2 around 300 ms in production and later, around 500 ms, in comprehension. Consistently, Strijkers and Costa (2016) suggest that top-down intentions (including the intention to speak) originating in the prefrontal cortex and in the temporo-parietal junction can influence the timing of subsequent computations of word production.

### 4.4 Limitations and future directions

Our study allows us to bring brain oscillatory evidence for the engagement of the production system in prediction during spoken language comprehension. However, our results do not allow us to make strong claims on the exact representations involved, specifically whether phonology is activated or not. Techniques with higher spatial resolution (such as MEG), experimental manipulations that better elicit phonological planning, and the study of special populations with speech–language disorders would contribute to characterize the cortical locations of the effects and help in understanding what representational levels are implicated. In addition, it must be underlined that EEG oscillatory activity is merely correlated with the observed experimental conditions. Because of this, we cannot make strong claims about whether the activity in a given brain area is necessary to a given process. However, the results of this study set the basis for further investigations with neurostimulation techniques (such as TMS) that could tease apart and clarify the role of these areas in language comprehension and production. Finally, individual differences may heavily hinder the ability to detect shared cognitive elaboration and neural activation, since we are assuming that if there are shared processes, they are unfolding at the same time in the two tasks; this may well not be the case. Despite these limitations, we argue that we bring sufficient evidence to stimulate further research along these lines, in an emerging effort to reconcile the study of language comprehension and production (McQueen & Meyer, 2019).

## 5. Conclusion

In this study we tested whether prediction-by-production accounts are supported by patterns of alpha and beta neural oscillations. Participants performed both a comprehension and a production task with predictable and non-predictable (but always plausible) target stimuli following constraining and non-constraining incomplete sentences. To our knowledge, this is the first attempt at studying both processes in the same set of participants, thereby investigating how the same mind–brain tackles the two tasks and directly comparing their neural responses. In addition, we employed naturalistic auditory stimuli differently from previous studies, replicating the modulations in a less artificial setting. We found alpha and beta power decrease (desynchronization) before predictable targets in both tasks, signaling that participants were retrieving and encoding rich linguistic information, compatible with the ‘information via desynchronization’ hypothesis. Source estimation and correlations suggest that 1) participants engage the left-lateralized word production network when predicting in comprehension and 2) internal modeling and contextual prediction is aided by the right temporo-parietal junction in both comprehension and production. These results stress the strict relationship between production and comprehension processes, lending support to prediction-by-production models.

## CRediT author statement

### Simone Gastaldon

Conceptualization, Methodology, Investigation, Formal analysis, Writing - Original Draft, Writing - Review & Editing, Visualization.

### Giorgio Arcara

Methodology, Software, Formal analysis, Writing - Review & Editing.

### Eduardo Navarrete

Conceptualization, Methodology, Writing - Review & Editing.

### Francesca Peressotti

Conceptualization, Methodology, Writing - Review & Editing, Supervision, Project administration

## Acknowledgments

SG was supported by a PhD grant from the University of Padova (2017–2020). GA was supported by the Italian Ministry of Health under Grant Number GR-2018-12366092. We thank all the participants for their collaboration and Bianca Bonato for recording the audio stimuli.

## Conflict of Interest Statement

The authors declare no competing interests.

## Appendix A. Supplementary data

Supplementary material for this manuscript can be found on the Open Science Framework at the following URL: https://osf.io/tcbsh/

## Footnotes

1 It is also worth considering that alpha and beta oscillations have been found to be implicated in a variety of functions, both outside and within the language domain, e.g. sensorimotor processing in action (Zaepffel, Brovelli, MacKay, and Riehle, 2013) and speech planning and execution (Saltuklaroglu et al., 2018), motor imagery and action semantics, working memory, information binding (Weiss & Mueller, 2012 for a review), time perception (Wiener, Parikh, Krakow, and Coslett, 2018), and temporal expectations (Morillon & Baillet, 2017).

2 If any channels were marked as ‘bad’, the number of components for the ICA was reduced to the number of good channels.

3 The delta band (0.1-4 Hz) was excluded because the wavelets at these frequencies were too large and temporal smearing introduced noise in the production task (HC condition) due to muscle activity after the gap of interest.

4 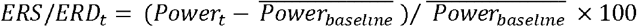

5 For the other layers (scalp, inner and outer skull) Brainstorm defaults settings were kept.

6 The other settings were kept at Brainstorm default settings (Noise covariance regularization: 0.1; Signal-to-noise ratio: 3).

## Notes

https://osf.io/tcbsh/

### Competing Interest Statement

The authors have declared no competing interest.

